# Mitochondrial Genome of *Garcinia mangostana* L. variety Mesta

**DOI:** 10.1101/2022.02.23.481586

**Authors:** Ching-Ching Wee, Nor Azlan Nor Muhammad, Vijay Kumar Subbiah, Masanori Arita, Yasukazu Nakamura, Hoe-Han Goh

## Abstract

Fruits of *Garcinia mangostana* L. (mangosteen) are rich in nutrients with xanthones found in the pericarp having great pharmaceutical potential. Mangosteen variety Mesta is only found in Malaysia, which tastes sweeter than the common Manggis variety in Southeast Asia. In this study, we report the complete mitogenome of *G. mangostana* L. variety Mesta with a total sequence length of 371,235 bp of which 1.7% could be of plastid origin. The overall GC content of the mitogenome is 43.8%, comprising 29 protein-coding genes, 3 rRNA genes, and 21 tRNA genes. Repeat and tandem repeat sequences accounted for 5.8% and 0.15% of the Mesta mitogenome, respectively. There are 333 predicted RNA-editing sites in Mesta mitogenome. These include the RNA-editing events that generated the start codon of *nad1* gene and the stop codon of *ccmFC* gene. Phylogenomic analysis with a maximum likelihood method showed that the mitogenome of mangosteen variety Mesta was grouped under Malpighiales order. This is the first complete mitogenome from the *Garcinia* genus for future evolutionary studies.

## Introduction

Mitochondria are the main organelle of energy production for cell sustainability. Besides chloroplast genomes (plastomes), mitochondrial genomes (mitogenomes) serve as important genetic information for phylogenetic and evolutionary studies^1^. The first land plant mitogenome sequenced was *Marchantia polymorpha*^2^. Mitogenome sizes are more variable, which range from 66 kb^3^ to 5.5 Mb^4^ compared to plastomes that are more conserved with the length range from 110 kb to 200 kb^5^. Plant mitogenomes are complex due to the rearrangement, duplication, recombination, and horizontal gene transfer between nucleus and organelles (plastids and mitochondria)^6-8^.

With the advancement of third-generation sequencing technology such as Pacific Biosciences (PacBio) and Oxford Nanopore Technologies (ONT) which can sequence over 100 kb per molecule and up to 1 million bp, long-read sequencing is becoming more commonly applied in mitogenome assembly^9,10^. The repetitive regions and rearrangement events that are common in mitogenomes hinder assembly with short-read sequencing^11^.

Mangosteen (*Garcinia mangostana* L.) is well-known as the “Queen of fruits” with sweet and juicy fruit pulp. It is a tropical fruit under the Clusiaceae family^12^ that can be found in Southeast Asia countries^13^. In Malaysia, there is a unique variety Mesta, which is characterized by an oblong shape, thicker mesocarp, and relatively fewer and smaller seeds compared to the common Manggis variety. Despite eight published chloroplast genomes from *Garcinia* species, there is no mitogenome from *Garcinia* species reported to date.

In this study, we assembled a complete mitogenome of Mesta using PacBio data^14^ and polished using Illumina short reads^15^. We also compared its structure and gene contents with five closely related species from the same order of Malpighiales and another two species from the Brassicales order, namely *Arabidopsis thaliana* and *Carica papaya* (reference used during assembly). The study provides a reference for future evolutionary studies of the *Garcinia* genus.

## Results

### General features of *Garcinia mangostana* var. Mesta mitogenome

*De novo* assembly using Organelle_PBA generated two mitogenome contigs with the length of 389,277 bp (scf7180000000010) and 20,340 bp (scf7180000000011), respectively. The smaller contig was the subset of the larger (master) contig (Supplementary Figure S1). For the larger contig that was circular, manual curation was done by removing one of the identical ends (∼18 kb) and a total of 63 bases were added based on the detected variants when short reads were aligned to the trimmed mitogenome (Supplementary Figure S2). The final complete mitogenome of *Garcinia mangostana* var. Mesta was 371,235 bp which was slightly larger than *Arabidopsis thaliana* mitogenome (367,808 bp) but smaller than *Carica papaya* mitogenome (476,890 bp) (Table 1). The average Mesta mitogenome coverage was 129 × using PacBio subreads (Supplementary Figure S3). The mitogenome comprising 29 protein-coding genes, 3 rRNA genes (*rrn5, rrn18*, and *rrn26*), and 21 tRNA genes (Figure 1 & Table 2). The total length of protein-coding genes was 28,113 bp, which accounted for 7.6% of the mitogenome. There were only five ribosomal proteins (*rpl5, rpl10, rpl16, rps3*, and *rps4*) found in the Mesta mitogenome.

**Table 1.** Comparison of the gene content in mitogenomes of different species.

| Order | Family | Features | Accession number | Size (bp) | GC content (%) | Number of gene | Protein-coding | rRNA | tRNA |
| --- | --- | --- | --- | --- | --- | --- | --- | --- | --- |
| Brassicales | Brassicaceae | *A. thaliana* | BK010421 | 367,808 | 44.8 | 58 | 33 | 3 | 22 |
|  | Caricaceae | *C. papaya* | NC_012116.1 | 476,890 | 45.1 | 61 | 39 | 3 | 19 |
| Malpighiales | Clusiaceae | *G. mangostana* | OM759996 | 371,235 | 43.8 | 55 | 29 | 3 | 21 |
|  | Euphorbiaceae | *R. communis* | NC_015141.1 | 502,773 | 45.0 | 60 | 37 | 3 | 20 |
|  | Salicaceae | *P. alba* | NC_041085.1 | 838,420 | 44.8 | 58 | 33 | 3 | 22 |
|  |  | *P. davidiana* | NC_035157.1 | 779,361 | 44.8 | 58 | 33 | 3 | 22 |
|  |  | *P. tremula* | NC_028096.1 | 783,442 | 44.7 | 58 | 33 | 3 | 22 |
|  | Passifloraceae | *P. edulis* | NC_050950.1 | 680,480 | 44.7 | 73 | 41 | 3 | 29 |

**Table 2.** List of genes in *Garcinia mangostana* var. Mesta mitogenome.

| Group | Gene |
| --- | --- |
| Complex I | *nad1, nad2, nad3, nad4, nad4L, nad5, nad6, nad7, nad9* |
| Complex III | *cob* |
| Complex IV | *cox1, cox2, cox3* |
| Complex V | *atp1, atp4, atp6, atp8, atp9* |
| Cytochrome c biogenesis | *ccmB, ccmC, ccmFc, ccmFn* |
| Ribosome large subunit | *rpl5, rpl10, rpl16* |
| Ribosome small subunit | *rps3, rps4,* |
| Intron maturase | *matR* |
| SecY-independent transporter | *mttB* |
| rRNA genes | *rrn5, rrnL, rrnS* |
| tRNA genes | *trnC-GCA, trnD-GUC, trnE-UUC, trnF-GAA (X2),*<br>*trnG-GCC, trnH-GUG, trnK-UUU, trnM-CAU (X5),*<br>*trnN-GUU (X2), trnP-UGG (X2), trnQ-UUG, trnS-*<br>*GCU, trnW-CCA, trnY-GUA* |

**Figure 1.**
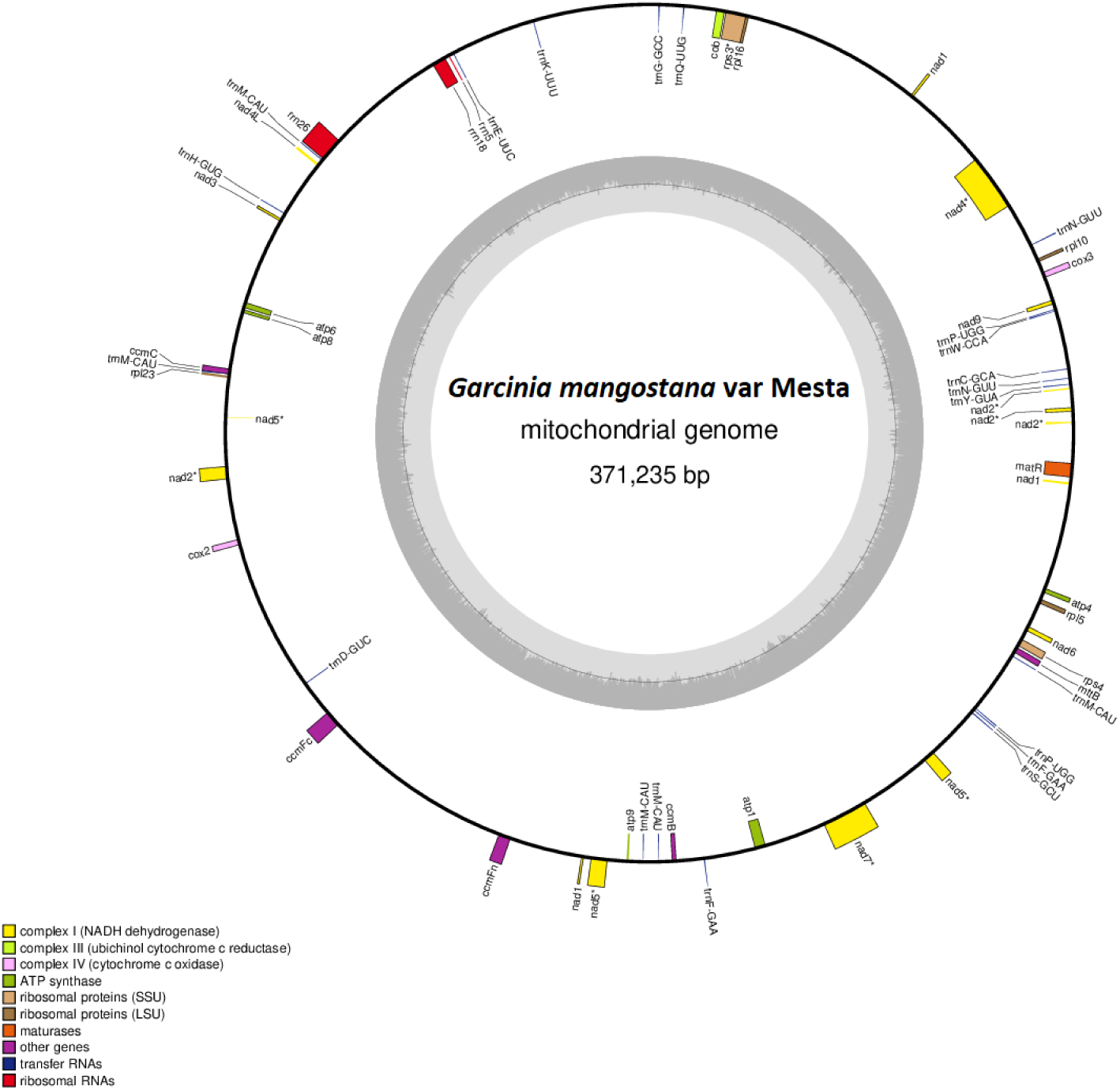
The circular mitochondrial genome of *G. mangostana* variety Mesta. Genes inside the circle are transcribed clockwise while genes outside the circle are transcribed anti-clockwise as indicated by the gray arrows. The gray bars inside the circle represent the GC content of the sequence. Asterisks (*) indicate genes containing intron(s).

**Figure 2.**
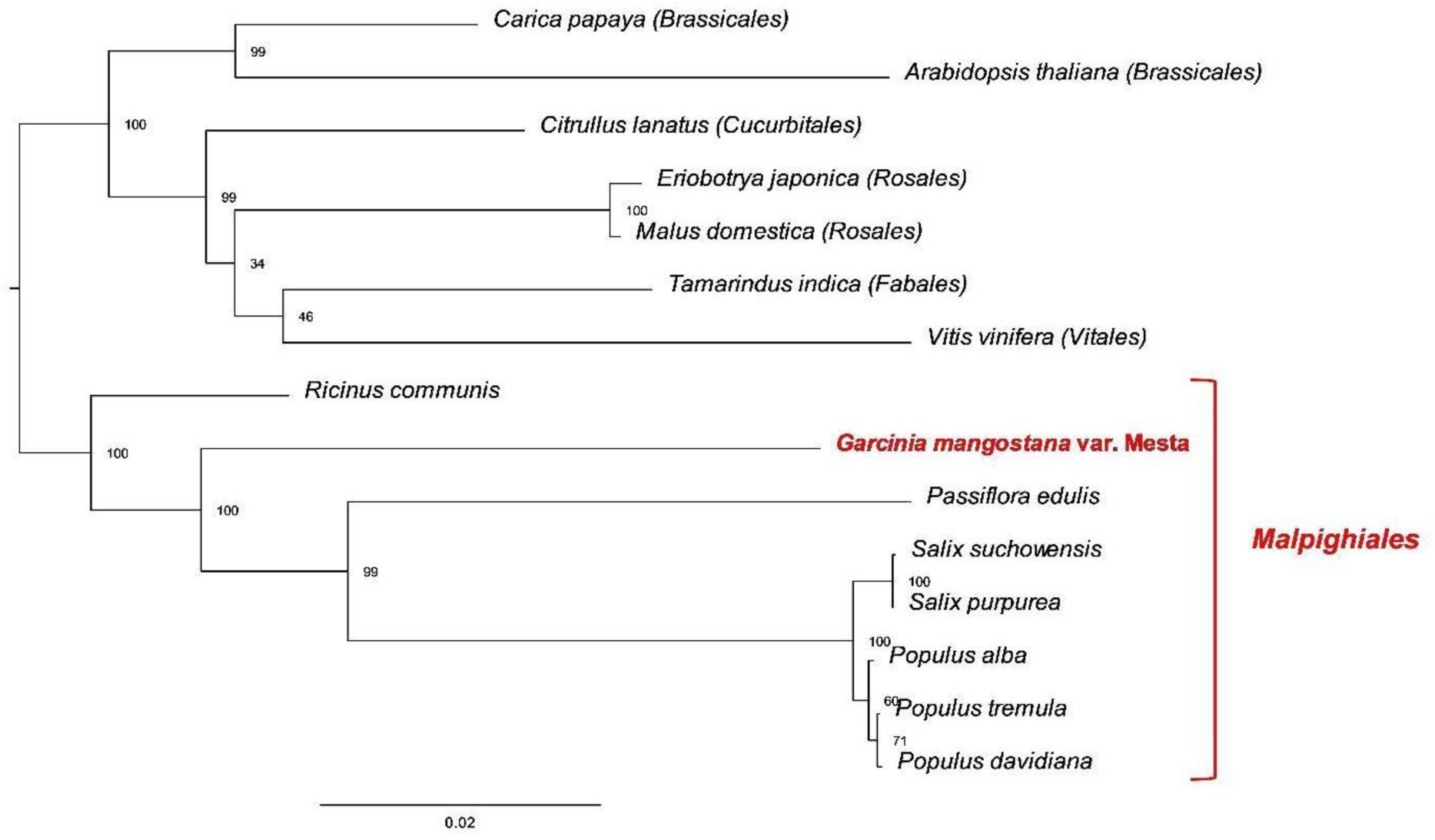
Phylogenomic tree (maximum likelihood) of 15 species based on 22 protein-coding genes. The parentheses and red line indicate the order name.

### Comparison of Mitogenome Gene content of different species

Comparison of the mitogenomes gene content of different species (Supplementary Table S1) showed that all the mitogenomes encoded the basic core set of 24 protein-coding genes. However, the mitogenome of *Passiflora edulis* encoded two copies of the genes *ccmB, nad4L, nad6*, and *nad7* and four copies of the gene *cox2*. Mitogenome of *Arabidopsis thaliana* also encoded two copies of *atp6* gene with the length of 1158 bp and 1050 bp, respectively. Two copies of *ccmFN* gene were also found in the mitogenome of *Carica papaya*. The *nad1* gene in Mesta was found only consisted of 3 exons instead of 5 as observed in other species despite the total length of its CDS sequence being almost like the other species. Most of the ribosomal proteins were found missing in both Salicaceae and Clusiaceae families from the order Malpighiales. Noticeably, gene *rps12* was missing from Mesta.

### Distribution of tRNAs

The 21 tRNAs identified in Mesta mitogenome only code for 14 amino acids (Ser, Phe, Asn, Met, Pro, Gly, Lys, Gln, Tyr, His, Trp, Asp, Glu, Cys). Two out of 21 tRNAs had a chloroplast-origin while the rest were mitochondrial-origin. However, the tRNA genes code for the other six amino acids (Leu, Ile, Thr, Ala, Val, Arg) were not detected. Among the 21 tRNAs, one of them was predicted to have one intron (Supplementary Table S2).

### Plastome-derived sequences

There were five plastome-derived sequences with an identity of more than 80% and a sequence length of at least 100 bp (Table 3) found in the mitogenome. This accounted for a total length of 6,214bp which was 1.7% of the mitogenome. The plastome genes contained were *rpl2* (partial), *rpl23, trnl-CAU, ndhA* (partial), *ndhH, rps15* (partial), *atpE* (partial), *atpB, rps3* (partial), and *trnD-GUC*. Three of the fragments were found at the plastome large single-copy (LSC) region and one from the single-copy (SSC) region and inverted repeat (IR) regions, respectively.

**Table 3.** Plastid gene insertions in the mitochondrial genome of Mesta.

| Plastid gene | Length (bp) | Position | Identity (%) | Location |
| --- | --- | --- | --- | --- |
| *rpl2*<sup>^</sup>, *rpl23*, *trnI-CAU* | 1,418 | 178,888-177,471 | 97.77 | IR |
| *ndhA*<sup>^</sup>, *ndhH*, *rps15*<sup>^</sup> | 2,476 | 21,314-18,839 | 97.68 | SSC |
| *atpE*<sup>^</sup>, *atpB* | 1,882 | 72,995-71,114 | 97.54 | LSC |
| *rps3*<sup>^</sup> | 289 | 310,031-310,319 | 87.06 | LSC |
| *trnD-GUC* | 149 | 222,409-222,557 | 85.81 | LSC |

<sup>^</sup> partial

### Introns and RNA Editing

There were 20 introns distributed among 8 protein-coding genes (*ccmFC, cox1, nad1, nad2, nad4, nad5, nad7*, and *rps3*) in Mesta mitogenome. Among them, *nad1, nad2*, and *nad5* genes were trans-spliced. The maximum number of introns found in a gene were four which can be found at certain *nad* gene families such as *nad2, nad5*, and *nad7*.

A total of 333 putative RNA-editing sites had been predicted using PREP-MT (Supplementary Table S4). Among the CDS, gene *nad4* contained the highest number of predicted editing sites (35) while there was no editing site found at the *atp9* gene. A total of three annotated genes did not start with the start codon ATG (*atp6, nad1*, and *rpl16*) (Supplementary Table S3). Among them, the ACG site at the beginning sequence of *nad1* was putatively predicted to be one of the editing sites that converted it into the ATG start codon. Similarly, early termination was predicted at the gene *ccmFC* sequence which converted the CGA into the stop codon TGA.

### Repeat sequences of Mesta mitochondrial DNA

A total of 64 repeat sequences comprising the forward and palindromic repeats were detected. The repeat sequences ranged from 51 bp to 4,172 bp with a total size of 21,356 bp which accounted for 5.8% of the total Mesta mitogenome (Supplementary Table S5). There were three forward repeat sequences with lengths of 4,172 bp, 2,031 bp, and 1,641 bp. As for the tandem repeats, it constituted 0.15% of the mitogenome (Supplementary Table S6).

### Phylogenomic Analysis

A total of 22 (excluding *ccmFn* and *mttB*) out of 24 basic core sets of protein-coding genes were used for phylogenomic analysis, which separated the 15 species into different groups based on the order. Mesta was grouped under Malpighiaes together with *P. edulis, S. suchowensis, S. purpurea, P. alba, P. tremula*, and *P. davidiana*.

## Discussion

Due to the high complexity of plant mitogenome with large repetitive regions, long-read sequencing is superior in mitogenome assembly^4,16^. In this study, a total of two Mesta mitogenome contigs were obtained using PacBio data. The shorter contig was a subset of the longer one with the size of 371,235 bp (after manual curation) and was considered as the complete Mesta mitogenome. It is not uncommon to have multiple mitogenome contigs in plants to exist in both circular and linear structures due to intramolecular recombination events^11,17,18^. For instance, there were 10 contigs in *Fagopyrum esculentum*^19^ and 13 contigs in *Picea sitchensis*^4^.

Mesta mitogenome encoded the basic core set of 24 protein-coding genes^20^ that include genes encoding proteins of the electron transport chain (complex I, III, IV, V, and cytoplasmic membrane proteins) commonly found in plant mitogenome^21^. However, Mesta mitogenome size was relatively small compared to other plant mitogenomes^22^ from the same order, Malpighiales (Table 1), due to the reduced number of ribosomal proteins and missing genes encoding respiratory chain complex II, *sdh3* and *sdh4* (Supplementary Table S3). These protein-coding genes could be lost during evolution and might be transferred to the nuclear genome as observed in other angiosperm mitogenomes^23-25^ such as *S. latifolia*^26^, *S. noctiflora*^27^, *P. dactylifera*, and *A. indica*^21,25^. For instance, *rps12* was not found in Mesta mitogenome as well as *Oenothera* and *Zostera marina*^28,29^.

A complete set of tRNAs coding for 20 amino acids is required for protein translation in plant mitogenomes. However, currently, there was no complete set of tRNA genes found in the mitogenome of angiosperm^30^. For the Mesta mitogenome, a total of six amino acids encoded by tRNAs (Leu, Ile, Thr, Ala, Val, Arg) were not detected and these tRNAs types were generally reported missing in angiosperm mitogenomes^30^. In the mitogenome of *S. latifolia*, the majority of the tRNAs were reported lost with only nine types of amino acids^26^. The loss of tRNAs can be either replaced by tRNAs from the chloroplast or nuclear genome^26,31^.

There are several factors attributed to the differences of plant mitogenome lengths, including the integration of nuclear and plastid genomes as well as the number and length of non-coding regions^20^. Chloroplast sequences can be found in plant mitogenomes as there were integration events during evolution. The integration of plastome sequences into mitogenomes can range from 1 to 12%^32^. For the Mesta mitogenome, the integration rate was 1.7% which was smaller compared to other plants such as watermelon (7.6%)^33^, *P. dactylifera* (10.3%)^21^, and *C. pepo* (11.6%)^34^.

The repetitive regions found in the intergenic regions of mitogenome were in variable types such as short repeats, tandem repeats, and long complex repeats^21,22,35,36^. The large repeats (>1 kb) might cause homologous recombination and eventually lead to the different configuration of the mitogenomes^37^. Apart from large repeats, both direct and inverse repeats also contribute to the subgenomic molecules^7^. Repeats detected in Mesta mitogenome using web-based REPuter was low (5.8%) compared to *C. melo* (42.7%)^38^, *V. vinifera* (6.8%)^39^, and *N. colorata* (48.89%)^40^ but higher than *P. dactylifera* (2.3%)^21^. Similarly, the tandem repeats detected in Mesta mitogenome were also low which was 0.15% compared to 0.33% in *P. dactylifera*^21^.

RNA-editing events are essential in plant development and stress response^41^. The most common RNA editing events in plant organelles (mitochondria and plastids) were the conversion of C-to-U^42^. RNA editing can lead to the start codon/stop codon generation, eliminate premature stop codon, change the splicing site, affect the RNA structure, and cause instability of RNAs^41^. It is predicted that RNA-editing events generated the start codon in *nad1* and the stop codon in *ccmFC* genes of Mesta mitogenome. The start codon of *nad1* gene in several species such as *A. alpina*^43^, *B. stricta*^44^, and *C. rubella*^45^ was also formed by RNA editing. On the other hand, stop codon prediction in *ccmFC* gene sequence had also been reported in *A. thaliana* and *C. bursa-pastoris*^1^. The GTG in *rpl16* might be a translation start codon as similar observations were found in maize, *Marchantia*, and *Petunia* mitogenomes^46^.

## Conclusion

The complete mitogenome of Mesta was successfully assembled. The Mesta mitogenome length was relatively smaller than other species in the same order due to the loss of the majority of ribosomal proteins and both *sdh* genes. Phylogenetic analysis based on the 22 protein-coding genes among the 15 selected species showed that Mesta was clustered within the Malpighiales order. The mitogenome can serve as a good reference to study the regulation of the mitogenome genes.

## Materials and Methods

### Mitochondrial Genome Assembly

Genome sequences of Mesta variety were obtained from the NCBI SRA database with the accession numbers SRX2718652 to SRX2718659 for PacBio long-read data (9.5 Gb)^14^ and SRX270978 for Illumina short reads (50.2 Gb)^15^. CANU v2.0^47^ was used to perform PacBio raw data correction and trimming using default parameters. Next, non-mitogenome reads were removed by sequence alignment of each read against a *Carica papaya* mitogenome (accession no. NC_012116.1) which was used as the reference genome. Then, *de novo* assembly was performed using Organelle_PBA software^48^. Manual curation was performed to obtain the complete Mesta mitogenome.

### Genome annotation

Gene annotation was conducted using both GeSeq^49^ and MITOFY web server^34^. Web-based tRNA-scan v2.0 server (http://lowelab.ucsc.edu/tRNAscan-SE/index.html)^50^ was used to annotate tRNA genes. The physical mitogenome map was generated using Organellar Genome DRAW (OGDRAW v1.3.1) program with default parameters^51^.

### Identification of plastome derived sequences

Plastome-derived sequences were identified by aligning plastome (accession number: MZ823408) and mitogenome of *G. mangostana* var. Mesta (accession number: OM759996) using NCBI-Nucleotide BLAST (BLASTN) webserver (https://blast.ncbi.nlm.nih.gov/Blast.cgi) with at least 80% sequence identity and alignment length greater than 100 bp.

### Analysis of RNA-Editing and Substitution Rate

Putative RNA-editing sites in protein-coding genes of Mesta mitogenome were predicted using PREP-mt web-based program (http://prep.unl.edu/)^52^. The cut-off value was set at 0.6 to obtain an accurate prediction.

### Analysis of Repetitive Sequences

The repeat sequences were identified using web-based REPuter (https://bibiserv.cebitec.uni-bielefeld.de/reputer/)^53^ with a minimum length of repeat size of 50 bp. Web-based tool Tandem Repeats Finder version 4.09^54^ (https://tandem.bu.edu/trf/trf.basic.submit.html) was used to identify tandem repeats in Mesta mt using default parameter.

### Phylogenomic Analysis

To examine the evolutionary relationship of *G. mangostana* var. Mesta, a maximum likelihood (ML) phylogenomic tree was inferred based on 22 protein-coding genes (Figure 1). The amino acids sequences of these 22 protein-coding genes were concatenated and aligned using MAFFT version 7 online tool (https://mafft.cbrc.jp/alignment/server/)^55^. The maximum likelihood (ML) analysis was performed using RAxML-NG v1.0.2 tool^56^ based on the selected model STMTREV+I+G4+F in the ModelTest-NG v0.1.6 ^57^.

## Data Availability Statement

The complete mitogenome sequence of *Garcinia mangostana* var. Mesta has been submitted to GenBank with the accession number OM759996.

## Acknowledgments

We would like to acknowledge the support of this research by Universiti Kebangsaan Malaysia (UKM) Research University grant AP-2012-018. The group is currently supported by UKM Research University grant DIP-2020-005 (H-HG) and NIG-JOINT grant 2021 (2A2021) (YN and H-HG), Japan.

## Author Contribution Statement

C.C.W. and H.H.G. conceived and planned the experiment. C.C.W. wrote the paper and analyzed the data. Y.N. contributed to the data analysis. H.H.G., N.A.N.M, V.K.S., M.A., and Y.N. reviewed and edited the manuscript. All authors read and approved the manuscript.

## Additional information

Supplementary Tables: Figure S1-S3

Supplementary Figures: Table S1-S6

