## Supplementary material for "Mitochondrial Genome of *Garcinia mangostana* L. variety Mesta": Supplementary Figures.docx

**Figure S1.** Schematic representation of large and small contigs of Mesta mitogenome.

**Figure S2.** Dot plot analysis showing scf7180000000010 as a circular contig.

**Figure S3**. Mesta mitogenome PacBio read depth.

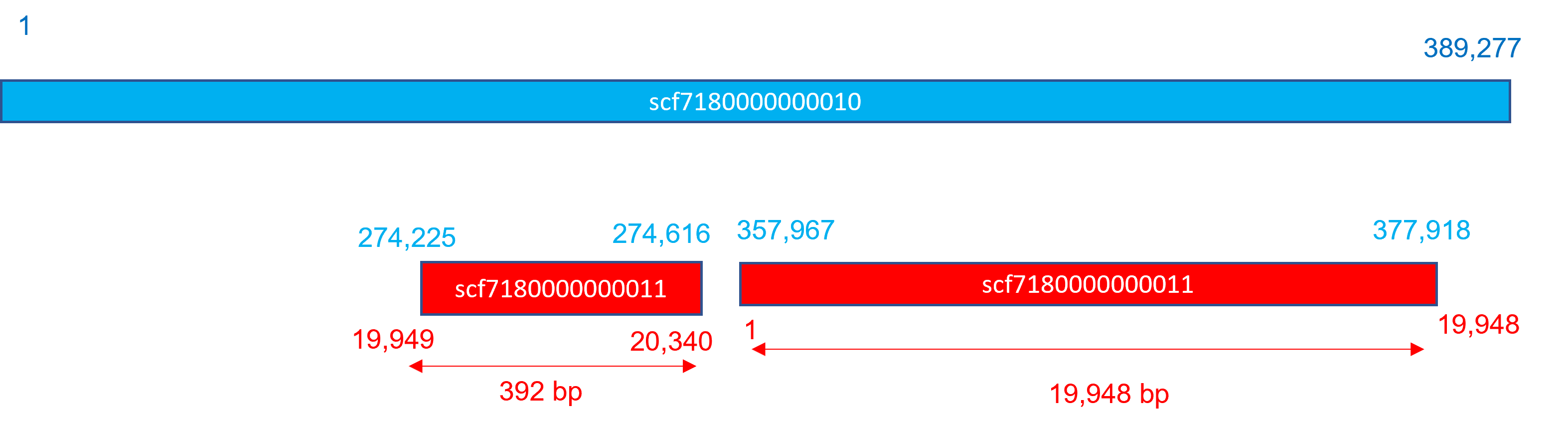

**Figure S1.** Schematic representation of large and small contigs of Mesta mitogenome.

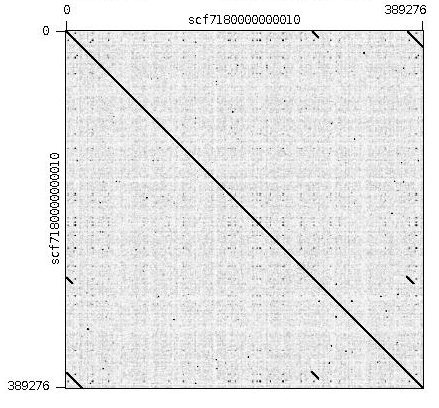

**Figure S2.** Dot plot analysis showing scf7180000000010 as a circular contig.

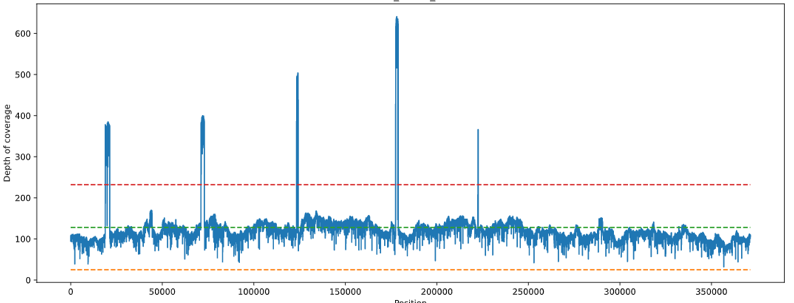

**Figure S3**. Mesta mitogenome PacBio read depth.
