## Supplementary material for "Mitochondrial Genome of *Garcinia mangostana* L. variety Mesta": Supplementary Tables.docx

**Table S1.** Length of protein-coding genes in mitogenomes of different species.

**Table S2.** tRNA gene content of *G. mangostana* mitogenome.

**Table S3.** Start codon and stop codon of the genes in Mesta mitogenome.

**Table S4.** Predicted RNA-editing sites in protein-coding genes of Mesta mitogenome.

**Table S5.** Repeats in the Mesta mitogenome.

**Table S6.** Tandem repeats in Mesta mitogenome.

**Table S1.** Length of protein-coding genes in mitogenomes of different species.

| **Gene** | ***A. thaliana*** | ***C. papaya*** | ***G. mangostana*** | ***R. communis*** | ***P. alba*** | ***P. davidiana*** | ***P. tremula*** | ***P. edulis*** |
| --- | --- | --- | --- | --- | --- | --- | --- | --- |
| *atp1* | 1524 | 1,530 | 1512 | 1,530 | 1524 | 1524 | 1524 | 1530 |
| *atp4* | 579 | 579 | 612 | 597 | 597 | 597 | 597 | 597 |
| *atp6* | 1158 | 774 | 768 | 792 | 714 | 714 | 714 | 1158 |
| *atp6*_2 | 1050 | - | - | - | - | - | - | - |
| *atp8* | 477 | 483 | 474 | 480 | 474 | 474 | 474 | 489 |
| *atp9* | 225 | 225 | 225 | 225 | 225 | 225 | 225 | 252 |
| *ccmB* | 621 | 621 | 621 | 621 | 615 | 615 | 615 | 621 (X2) |
| *ccmC* | 771 | 726 | 753 | 699 | 753 | 753 | 753 | 753 |
| *ccmFC* exon1 | 779 | 773 | 762 | 767 | 767 | 767 | 767 | 767 |
| *ccmFC* exon*2* | 550 | 550 | 549 | 550 | 595 | 595 | 595 | 544 |
| *ccmFN* | 1149 | 1731 (X2) | 1722 | 1,797 | 1731 | 1731 | 1731 | 1749 |
| *ccmFN2* | Ψ | - | - | - | - | - | - | - |
| *cob* | 1182 | 1,182 | 1182 | 1,182 | 1182 | 1182 | 1182 | 1173 |
| *cox1* exon1 | 1584 | 1,584 | 717 | 726 | 1584 | 1584 | 1584 | 1833 |
| *cox1* exon2 | - | - | 858 | 858 | - | - | - | - |
| *cox2* exon1 | 700 | 699 | 795 | 700 | 675 | 675 | 675 | 777 (X2) |
| *cox2* exon2 | 83 | 84 | - | 83 | - | - | - | - |
| *cox2*_2 | - | - | - | - | - | - | - | 1257 (X2) |
| *cox3* | 798 | 798 | 798 | 798 | 798 | 798 | 798 | 798 |
| *matR* | 1971 | 1,968 | 1959 | 1,968 | 1944 | 1944 | 1944 | 1971 |
| *mttB* | 843 | Ψ | Ψ | Ψ | - | Ψ | Ψ | 852 |
| *nad1* exon1 | 385 | 385 | 387 | 387 | 298 | 298 | 298 | 297 |
| *nad1* exon2 | 83 | 83 | 332 | 81 | 80 | 80 | 80 | 81 |
| *nad1* exon3 | 192 | 192 | 259 | 192 | 192 | 192 | 192 | 252 |
| *nad1* exon4 | 59 | 59 | - | 59 | 59 | 59 | 59 | 258 |
| *nad1* exon5 | 259 | 259 | - | 259 | 259 | 259 | 259 | - |
| *nad2* exon1 | 153 | 153 | 153 | 153 | 153 | 153 | 153 | 153 |
| *nad2* exon2 | 392 | 392 | 392 | 392 | 392 | 392 | 392 | 393 |
| *nad2* exon3 | 161 | 161 | 161 | 161 | 155 | 155 | 155 | 162 |
| *nad2* exon4 | 573 | 573 | 572 | 573 | 573 | 573 | 573 | 561 |
| *nad2* exon5 | 188 | 188 | 189 | 188 | 188 | 188 | 188 | 198 |
| *nad3* | 357 | 357 | 357 | 357 | 357 | 357 | 357 | 357 |
| *nad4* exon1 | 461 | 461 | 461 | 461 | 461 | 461 | 461 | 461 |
| *nad4* exon2 | 515 | 515 | 515 | 515 | 515 | 515 | 515 | 517 |
| *nad4* exon3 | 423 | 423 | 422 | 423 | 423 | 423 | 423 | 421 |
| *nad4* exon4 | 89 | 89 | 90 | 89 | 89 | 89 | 89 | 89 |
| *nad4L* | 303 | 303 | 303 | 303 | 303 | 303 | 303 | 303 (X2) |
| *nad5* exon1 | 230 | 230 | 230 | 230 | 229 | 230 | 230 | 229 |
| *nad5* exon2 | 1216 | 1,216 | 1216 | 1,216 | 1218 | 1216 | 1216 | 1217 |
| *nad5* exon3 | 22 | 22 | 22 | 22 | 21 | 22 | 22 | 396 |
| *nad5* exon4 | 395 | 395 | 395 | 395 | 395 | 395 | 395 | 150 |
| *nad5* exon5 | 147 | 147 | 150 | 150 | 138 | 138 | 138 | - |
| *nad6* | 618 | 618 | 618 | 654 | 630 | 630 | 630 | 615 (X2) |
| *nad7* exon1 | 143 | 143 | 143 | 143 | 143 | 143 | 143 | 143 (X2) |
| *nad7* exon2 | 69 | 69 | 69 | 69 | 69 | 69 | 69 | 69 (X2) |
| *nad7* exon3 | 467 | 467 | 467 | 467 | 467 | 467 | 467 | 467 (X2) |
| *nad7* exon4 | 244 | 244 | 244 | 244 | 244 | 244 | 244 | 244 (X2) |
| *nad7* exon5 | 262 | 262 | 262 | 262 | 262 | 262 | 262 | 262 (X2) |
| *nad9* | 573 | 573 | 573 | 573 | 573 | 573 | 573 | 573 |
| *rpl2* exon1 | 917 | 890 | - | - | 891 | 891 | 891 | - |
| *rpl2* exon2 | 133 | 118 | - | - | 117 | 117 | 117 | - |
| *rpl5* | 558 | 558 | 615 | 555 | - | - | - | 561 |
| *rpl10* | - | - | 441 | 453 | 489 | 489 | 489 | 465 |
| *rpl16* | 540 | 435 | 447 | 435 | Ψ | - | - | 249 |
| *rps1* exon1 | - | 603 | - | 606 | 636 | 636 | 636 | 226 |
| *rps1* exon2 | - | - | - | - | - | - | - | 443 |
| *rps3* exon1 | 74 | 74 | 74 | 74 | 69 | 69 | 69 | 74 |
| *rps3* exon2 | 1597 | 1,579 | 1411 | 1618 | 1575 | 1575 | 1575 | 1615 |
| *rps4* | 1089 | Ψ | 1026 | 1,059 | 969 | 969 | 969 | 954 (X2) |
| *rps7* | 447 | 450 | - | 447 | 447 | 447 | 447 | - |
| *rps10* exon1 | - | 250 | - | 250 | - | - | - | - |
| *rps10* exon2 | - | 83 | - | 83 | - | - | - | - |
| *rps12* | 378 | 378 | - | 378 | 378 | 378 | 378 | 378 |
| *rps13* | - | 351 | - | 351 | - | - | - | - |
| *rps14* | - | 303 | - | - | 246 | 246 | 246 | - |
| *rps19* | - | 285 | - | 195 | - | - | - | 285 |
| *sdh3* | - | 327 | - | 312 | - | - | - | 288 |
| *sdh4* | - | Ψ | - | 393 | 396 | 396 | 396 | - |

Ψ = pseudogene or fragment

**Table S2.** tRNA gene content of *G. mangostana* mitogenome.

| **tRNA type** | **tRNA Begin** | **tRNA End** | **tRNA Type** | **Intron Begin** | **Intron End** | **Length (bp)** |
| --- | --- | --- | --- | --- | --- | --- |
| *trnC-GCA* | 8967 | 8897 | Cys | - | - | 71 |
| *trnD-GTC* | 222555 | 222482 | Asp | - | - | 74 |
| *trnE-TTC* | 121253 | 121182 | Glu | - | - | 72 |
| *trnF-GAA* | 286030 | 286109 | Phe | - | - | 80 |
| *trnF-GAA* | 329797 | 329870 | Phe | - | - | 74 |
| *trnG-GCC* | 91506 | 91428 | Gly | - | - | 79 |
| *trnH-GTG* | 153864 | 153937 | His | - | - | 74 |
| *trnK-TTT* | 109241 | 109169 | Lys | - | - | 73 |
| *trnM-CAT* | 144816 | 144888 | Met | - | - | 73 |
| *trnM-CAT* | 177471 | 177544 | Met | - | - | 74 |
| *trnM-CAT* | 338961 | 339033 | Met | - | - | 73 |
| *trnM-CAT* | 279775 | 279702 | Met | - | - | 74 |
| *trnM-CAT* | 277570 | 277498 | Met | - | - | 73 |
| *trnN-GTT* | 26757 | 26843 | Asn | - | - | 87 |
| *trnN-GTT* | 7659 | 7588 | Asn | - | - | 72 |
| *trnP-TGG* | 330108 | 330182 | Pro | - | - | 75 |
| *trnP-TGG* | 17349 | 17272 | Pro | 17311 | 17308 | 78 |
| *trnQ-TTG* | 88028 | 87957 | Gln | - | - | 72 |
| *trnS-GCT* | 329347 | 329438 | Ser | - | - | 92 |
| *trnW-CCA* | 17136 | 17063 | Trp | - | - | 74 |
| *trnY-GTA* | 6852 | 6770 | Tyr | - | - | 83 |

**Table S3.** Start codon and stop codon of the genes in Mesta mitogenome.

| **Gene name** | **Length (bp)** | **Start codon** | **Stop codon** |
| --- | --- | --- | --- |
| *atp1* | 1512 | ATG | TAA |
| *atp4* | 612 | ATG | TAG |
| *atp6* | 768 | ACG | TAA |
| *atp8* | 474 | ATG | TAA |
| *atp9* | 225 | ATG | TAA |
| *ccmB* | 621 | ATG | TGA |
| *ccmC* | 753 | ATG | TGA |
| *ccmFC* | 1311 | ATG | CGA |
| *ccmFN* | 1722 | ATG | TGA |
| *cob* | 1182 | ATG | TAA |
| *cox1* | 1575 | ATG | TAA |
| *cox2* | 795 | ATG | TAA |
| *cox3* | 798 | ATG | TGA |
| *matR* | 1959 | ATG | TAG |
| *mttB* | 780 | Not determined | TAG |
| *nad1* | 978 | ACG | TAA |
| *nad2* | 1467 | ATG | TAA |
| *nad3* | 357 | ATG | TAA |
| *nad4* | 1488 | ATG | TGA |
| *nad4L* | 303 | ATG | TAA |
| *nad5* | 2013 | ATG | TAA |
| *nad6* | 618 | ATG | TAA |
| *nad7* | 1185 | ATG | TAG |
| *nad9* | 573 | ATG | TAA |
| *rpl5* | 615 | ATG | TAG |
| *rpl10* | 441 | ATG | TAA |
| *rpl16* | 447 | GTG | TAA |
| *rps3* | 1485 | ATG | TAA |
| *rps4* | 1026 | ATG | TAA |

**Table S4.** Predicted RNA-editing sites in protein-coding genes of Mesta mitogenome.

| **Gene** | **Number of predicted RNA-editing sites** |
| --- | --- |
| *atp1* | 3 |
| *atp4* | 7 |
| *atp6* | 6 |
| *atp8* | 3 |
| *atp9* | 0 |
| *ccmB* | 30 |
| *ccmC* | 25 |
| *ccmFC* | 14 |
| *ccmFN* | 27 |
| *cob* | 5 |
| *cox1* | 4 |
| *cox2* | 8 |
| *cox3* | 6 |
| *matR* | 10 |
| *mttB* | 22 |
| *nad1* | 13 |
| *nad2* | 21 |
| *nad3* | 4 |
| *nad4* | 35 |
| *nad4L* | 9 |
| *nad5* | 19 |
| *nad6* | 8 |
| *nad7* | 17 |
| *nad9* | 8 |
| *rpl5* | 6 |
| *rpl10* | 3 |
| *rpl16* | 4 |
| *rps3* | 6 |
| *rps4* | 10 |

**Table S5.** Repeats in the Mesta mitogenome.

| **Repeat length of the first fragment** | **Starting position of the first fragment** | **Repeat type** | **Repeat length of the second fragment** | **Starting position of the second fragment** | **E-value** |
| --- | --- | --- | --- | --- | --- |
| 4172 | 250350 | F | 4172 | 354064 | 0.00E+00 |
| 2031 | 248322 | F | 2031 | 352037 | 0.00E+00 |
| 1641 | 254519 | F | 1641 | 358232 | 0.00E+00 |
| 236 | 267849 | F | 236 | 342772 | 3.18E-132 |
| 221 | 270120 | F | 221 | 323741 | 3.41E-123 |
| 220 | 5096 | P | 220 | 305629 | 1.37E-122 |
| 183 | 275226 | F | 183 | 292371 | 2.58E-100 |
| 171 | 256809 | P | 171 | 275414 | 4.33E-93 |
| 130 | 179709 | P | 130 | 336133 | 2.09E-68 |
| 123 | 98918 | F | 123 | 295354 | 3.43E-64 |
| 117 | 36176 | P | 117 | 144975 | 1.40E-60 |
| 91 | 82130 | P | 91 | 270039 | 6.32E-45 |
| 91 | 285961 | F | 91 | 329727 | 6.32E-45 |
| 89 | 57528 | P | 89 | 219306 | 1.01E-43 |
| 86 | 145510 | F | 86 | 348707 | 6.47E-42 |
| 83 | 21838 | F | 83 | 348678 | 4.14E-40 |
| 82 | 124528 | P | 82 | 346426 | 1.66E-39 |
| 80 | 18686 | F | 80 | 168090 | 2.65E-38 |
| 79 | 320418 | F | 79 | 341651 | 1.06E-37 |
| 68 | 83466 | P | 68 | 108531 | 4.45E-31 |
| 66 | 3987 | F | 66 | 266272 | 7.12E-30 |
| 62 | 25035 | P | 62 | 275348 | 1.82E-27 |
| 61 | 25036 | P | 61 | 292493 | 7.29E-27 |
| 61 | 144883 | F | 61 | 339028 | 7.29E-27 |
| 59 | 86819 | F | 59 | 161722 | 1.17E-25 |
| 56 | 8143 | P | 56 | 364675 | 7.47E-24 |
| 56 | 18748 | P | 56 | 199898 | 7.47E-24 |
| 54 | 21867 | F | 54 | 145510 | 1.19E-22 |
| 54 | 152474 | F | 54 | 159083 | 1.19E-22 |
| 53 | 220661 | F | 53 | 361249 | 4.78E-22 |
| 51 | 76162 | F | 51 | 122259 | 7.64E-21 |
| 51 | 98908 | F | 51 | 168792 | 7.64E-21 |

**Table S6.** Tandem repeats in Mesta mitogenome.

| **Position** | **Repeat unit** | **Copy number** |
| --- | --- | --- |
| [21306--21390](https://tandem.bu.edu/trf/output/45BECvJ7ZP0us.2.7.7.80.10.50.500.1.txt.html#21306--21390,40,2.1,40,1) | 40 | 2.1 |
| [102358--102405](https://tandem.bu.edu/trf/output/45BECvJ7ZP0us.2.7.7.80.10.50.500.1.txt.html#102358--102405,24,2.0,24,2) | 24 | 2.0 |
| [148805--148841](https://tandem.bu.edu/trf/output/45BECvJ7ZP0us.2.7.7.80.10.50.500.1.txt.html#148805--148841,19,1.9,19,3) | 19 | 1.9 |
| [248585--248632](https://tandem.bu.edu/trf/output/45BECvJ7ZP0us.2.7.7.80.10.50.500.1.txt.html#248585--248632,21,2.3,21,4) | 21 | 2.3 |
| [250643--250723](https://tandem.bu.edu/trf/output/45BECvJ7ZP0us.2.7.7.80.10.50.500.1.txt.html#250643--250723,35,2.3,35,5) | 35 | 2.3 |
| [318648--318769](https://tandem.bu.edu/trf/output/45BECvJ7ZP0us.2.7.7.80.10.50.500.1.txt.html#318648--318769,65,1.9,65,6) | 65 | 1.9 |
| [352300--352347](https://tandem.bu.edu/trf/output/45BECvJ7ZP0us.2.7.7.80.10.50.500.1.txt.html#352300--352347,21,2.3,21,7) | 21 | 2.3 |
| [354357--354437](https://tandem.bu.edu/trf/output/45BECvJ7ZP0us.2.7.7.80.10.50.500.1.txt.html#354357--354437,35,2.3,35,8) | 35 | 2.3 |
